# Constrained enumeration of *k*-mers from a collection of references with metadata

**DOI:** 10.1101/2024.05.26.595967

**Authors:** Florian Ingels, Igor Martayan, Mikaël Salson, Camille Marchet

## Abstract

While recent developments in *k*-mers indexing methods have opened up many new possibilities, they still have limitations in handling certain types of queries, such as identifying *k*-mers present in one dataset but absent in another. In this article, we present a framework for efficiently enumerating all *k*-mers within a collection of references that satisfy constraints related to their metadata tags. Our method involves simplifying the query beforehand to reduce computation delays; the construction of the solution itself is carried out using CBL, a recent data structure specifically dedicated to the optimised computation of set operations on *k*-mer sets. We provide an implementation to our solution and we demonstrate its capabilities using real genomic data (microbial and RNA-seq), and show examples of use cases to identify *k*-mers of biological interest.

**Funding:** This work is funded by a grant from the French ANR: Full-RNA ANR-22-CE45-0007. Igor Martayan is supported by a doctoral grant from ENS Rennes.

## 1 Introduction

A few years ago the feasibility of indexing comprehensive genomic or transcriptomic datasets seemed beyond reach. However, recent advances in the indexing of large-scale biological sequence datasets have brought solutions leading to different large scale queryable indexes. Examples include the entirety of assembled bacterial genomes [5], or creating a searchable, quantitative index for RNA-seq data in cancer research [1]. Various methodologies, all of them based on *k*-mers, have been introduced to this extent. These methods typically represent *k*-mer sets in compact, compressed formats that also retain information about the origin of the *k*-mers. These indexing solutions often employ structures like colored de Bruijn graphs, among others [9].

In essence, these structures operate by breaking down a query sequence into *k*-mers and by identifying intersection between the *k*-mer set of the query and those of the indexed datasets. While these developments have unlocked opportunities, the range of queries that can be effectively handled remains limited. For example, current methods do not support inverse operations such as determining which *k*-mers are present in certain datasets but absent in others. This capability would be particularly useful in comparative studies, such as differentiating between datasets derived from patients versus control groups. This type of query would be more typical in databases. However, databases of *k*-mers are underdeveloped and notably prohibitive in ingestion time.

In this paper, we introduce a new framework designed to efficiently enumerate *k*-mers from various datasets based on specified constraints or metadata tags. We demonstrate how to construct subsets of *k*-mers using these tags and how to simplify query operations to reduce computational delays. We implement a solution using a recent *k*-mer set data-structure that was tailored for set operations [11]. We demonstrate that our method is capable of handling thousands of datasets efficiently. To illustrate its potential applications, we present two specific case studies showcasing the type of research that can be conducted using our approach.

## 2 Problem formulation

### 2.1 References and tags

Let ℛ = {*R*_1_, …, *R*_*N*_} be a collection of references (e.g. genomes, transcriptomes, etc.). A reference *R*_*i*_ is a string over DNA alphabet Σ = {*A, C, G, T*}. A *k*-mer is a word on Σ^*k*^, with *k* ≥ 1 fixed. Using the notion of *spectrum* from [12], we denote by *σ*(*R*) the spectrum of a reference *R*, i.e. the set of all *k*-mers of *R*: *σ*(*R*) = {*x* ∈ Σ^*k*^ : *x* ∈ *R*} – where *x* ∈ *R* is to be understood as “*x* is a substring of *R*”.

The spectrum of the collection ℛ is defined as

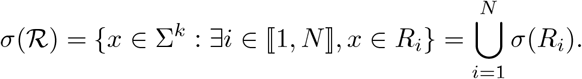

Since we assume the collection ℛ to be fixed in the sequel of this paper, and since *σ*(ℛ) denotes the set of all relevant *k*-mers, we shall also denote this set by 𝒰 and refer to it as “the universe”.

A metadata associated to a reference, is a boolean stating whether the reference has a given property. Thus each reference is associated with a list of *M* booleans {*t*_1_, …, *t*_*M*_} identifying the metadata they have. All the references with a given property (say the *i*-th one) are called a tag.

#### ▸ Definition 1.

*Any subset Q* ⊆ {1, …, *N*} *is referred to as a* tag.

Tags can be labeled with strings but, for the sake of simplicity, we will use capital letters *A, B, C*, … to designate tags in this article.

### 2.2 Building *k*-mers sets from tags

We want to identify *k*-mers that appear in references with some tags and that do not appear in references with some other tags. More formally, we consider here four operators All, Any, NotAll and NotAny, which can be applied to a tag *Q* as follows to construct sets of *k*-mers :

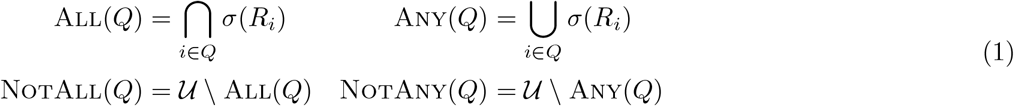

In other words, All(*Q*) represents the set of *k*-mers which appear in *all* the references of *Q*, whereas Any(*Q*) represents the set of *k*-mers which appear in *at least* one reference of *Q*. NotAny(*Q*) represents the set of *k*-mers which do not appear in *any* reference of *Q*; whereas NotAll(*Q*) represents the set of *k*-mers which do not appear *simultaneously* in all the references of *Q*.

The problem we are interested in this document is the following.

#### ▸ Problem 1.

*Provided four sets of tags* 𝒜 = {*A*_1_, *A*_2_, …}, ℬ = {*B*_1_, *B*_2_, …}, 𝒞 = {*C*_1_, *C*_2_, …} *et* 𝒟 = {*D*_1_, *D*_2_ …}, *build the following k-mer set:*

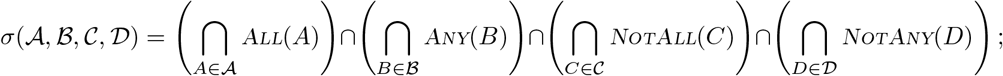

*with the understanding that if one of the sets* 𝒜, ℬ, 𝒞 *or* 𝒟 *is empty, the associated intersection is removed; by convention σ*(∅, ∅, ∅, ∅) = ∅.

In other words, *σ*(𝒜, ℬ, 𝒞, 𝒟) represents the set of all *k*-mers in 𝒰 that satisfy simultaneously all the required constraints. The quadruplet (𝒜, ℬ, 𝒞, 𝒟) is referred to as *query* in the following.

Throughout this article, we are going to propose a number of different reorganisations (using the usual set operations) of the way in which we write *σ*(𝒜, ℬ, 𝒞, 𝒟), with the aim of simplifying its writing and calculation. We start by the following one.

#### ▸ Proposition 2.

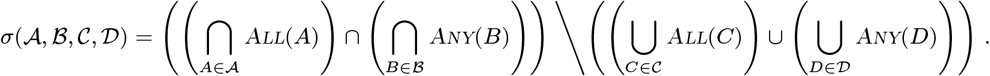

Proof. The proof derives naturally from the following manipulations, where *P* and *Q* are sets belonging to the same universe 𝒰 : (𝒰 \ *P*) ∩ (𝒰 \ *Q*) = 𝒰\ (*P* ∪ *Q*) and *P* ∩ ((𝒰 \ *Q*) = (*P* ∩ (𝒰)\*Q* = *P \Q*. ◂

With regard to Proposition 2, whenever 𝒜 = ℬ = ∅, the left part of \ should be replaced by 𝒰; whereas whenever 𝒞= 𝒟 = ∅, the right part of \ should be replaced by ∅. If all of them are empty sets, the solution set is also empty as per the convention of Problem 1.

## 3 Simplifying queries

Although we expect the majority of queries to be fairly straightforward, our method is intended to be general and allows for potentially complex queries. In particular, by manipulating only tags and forgetting, so to speak, the precise references associated with them, it is possible to construct queries almost in natural language, which we believe is convenient for the user. However, these queries, once translated into our framework, can turn out to be more complex than necessary in terms of the number of operations required to construct the solution. In this section, we therefore propose a method for simplifying the queries, leading to the same final set of *k*-mers but reducing, whenever possible, the quantity of operations required.

First, let us define the size of a query (𝒜, ℬ, 𝒞, 𝒟) as the quantity

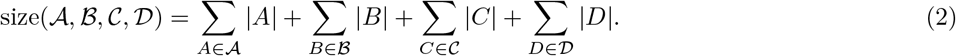

### ▸ Definition 3.

*We say that the query* (𝒜, ℬ, 𝒞, 𝒟) *simplifies into the query* (𝒜^*′*^, ℬ^*′*^, 𝒞^*′*^, 𝒟^*′*^) *if and only if* :

▄ *they induce the same k-mer set, i*.*e. σ*(𝒜^*′*^, ℬ^*′*^, 𝒞^*′*^, 𝒟^*′*^) = *σ*(𝒜, ℬ, 𝒞, 𝒟),
▄ *the size of the query decreases, i*.*e. size*(𝒜^*′*^, ℬ^*′*^, 𝒞^*′*^, 𝒟^*′*^) ≤ *size*(𝒜, ℬ, 𝒞, 𝒟).

The objective of this section is to simplify as much as possible the initial query (𝒜, ℬ, 𝒞, 𝒟).

### 3.1 𝒜 and 𝒟

We start by considering the following result.

#### ▸ Proposition 4.

*Let P and Q be two tags*.*Then, we have:*

▄ *All*(*P*) ∩ *All*(*Q*) = *All*(*P* ∪ *Q*),
▄ *Any*(*P*) ∪ *Any*(*Q*) = *Any*(*P* ∪ *Q*).

Proof. It follows trivially from the definition of operators All and Any – see (1). ◂

By denoting *A*^∪^ = ∪_*A*∈𝒜_ *A* and *D*^∪^ = ∪_*D*∈𝒟_ *D*, we deduce from Proposition 2 the following simplification of the query – where the decrease in the size of the query is immediate.

#### ▸ Corollary 5.

(𝒜, ℬ, 𝒞, 𝒟) *simplifies into* ({*A*^∪^}, ℬ, 𝒞, {*D*^∪^}). *Therefore, we have:*

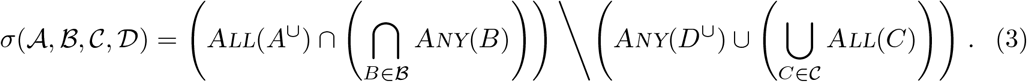

### 3.2 *A*^∪^ ∩ *D*^∪^, ℬ and 𝒞

Consider the following result.

#### ▸ Proposition 6.

*Let P, Q be tags so that P* ∩ *Q ≠* ∅. *Then, we have:*

i. *All*(*Q*) \ *Any*(*P*) = ∅,
ii. *All*(*Q*) ∩ *Any*(*P*) = *All*(*Q*),
iii. *All*(*Q*) ∪ *Any*(*P*) = *Any*(*P*), *Let us denote I* = *P* ∩ *Q, Q*^∗^ = *Q\ I and P* ^∗^ = *P\ I. Then, we also have:*
iv. *All*(*Q*) \ *All*(*P*) = *All*(*Q*) \ *All*(*P* ^∗^),
v. *Any*(*Q*) \ *Any*(*P*) = *Any*(*Q*^∗^) \ *Any*(*P*).

**Proof**. The proof is deferred to Supplementary, Section A.1. ◂

Note that, provided sets *P*_1_, … *P*_*n*_ and *Q*_1_, …, *Q*_*m*_, we have (*P*_1_ ∩ · · · ∩ *P*_*n*_) \(*Q*_1_ ∪ · · · ∪ *Q*_*m*_) = ∩_*i*,*j*_ (*P*_*i*_\*Q*_*j*_); hence, we can rewrite *σ* (𝒜, ℬ, 𝒞, 𝒟) as:

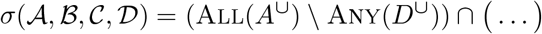

From Proposition 2–(i), we immediately conclude the following.

#### ▸ Corollary 7.

*If A*^∪^ ∩ *D*^∪^ ≠ ∅, *then σ*(𝒜, ℬ, 𝒞, 𝒟) = ∅.

In this case, we stop. Otherwise, assume that *A*^∪^ ∩ *D*^∪^ = ∅ and let us resume the simplification. Let *P, Q, R* and *S* be four sets, notice that (*P* ∩ *Q*)\ (*R* ∪ *S*) = (*Q\R*) ∩ (*P\S*). Consider again (3); we have:

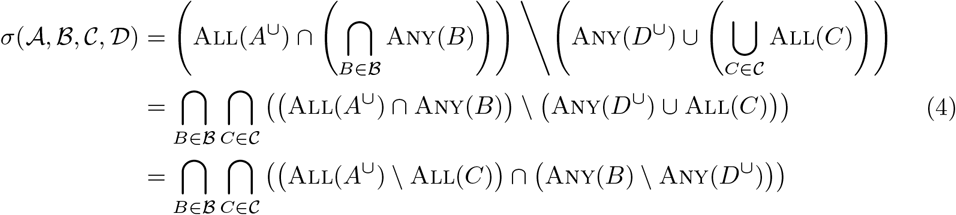

With regard to Proposition 2, one can notice the following facts:

▄ Combining second line of (4) with items (ii) and (iii) of Proposition 2 allows us to exclude from the calculation the sets *B* ∈ ℬ such that *B* ∩ *A*^∪^ ≠ ∅, as wells as for the sets *C* ∈ 𝒞 such that *C* ∩ *D*^∪^ ≠ ∅;
▄ Combining third line of (4) with items (iv) and (v) of Proposition 2 allows us to replace the sets *B* ∈ ℬ by *B \ D*^∪^, as well as the sets *C* ∈ 𝒞 by *C\A*^∪^.

The following result has therefore been proved. Once again, it is straightforward to see that the size of the query is reduced after these modifications.

#### ▸ Corollary 8.

(𝒜, ℬ, 𝒞, 𝒟) *simplifies into* ({*A*^∪^}, ℬ^∗^, 𝒞^∗^, {*D*^∪^}), *where* ℬ^∗^ = {*B\ D*^∪^ : *B* ∈ *B such that* ℬ ∩ *A*^∪^ = ∅} *and* 𝒞^∗^ = {*C \ A*^∪^ : *C* ∈ 𝒞 *such that C* ∩ *D*^∪^ = ∅}.

### 3.3 ℬ^∗^ and 𝒞^∗^

For this last round of simplifications, we start by the following result.

#### ▸ Proposition 9.

*Let P, Q be tags so that P* ⊆ *Q. Then, we have* :

i. *Any*(*P*) ∩ *Any*(*Q*) = *Any*(*P*),
ii. *All*(*P*) ∪ *All*(*Q*) = *All*(*P*).

**Proof**. Let us denote *Q*^∗^ = *Q* \ *P*. (i) It suffices to note that Any(*Q*) = Any(*P*) ∪ Any(*Q*^∗^) ⊇ Any(*P*) – hence, intersected against Any(*P*), we obtain Any(*P*). (ii) Similarly, All(*Q*) = All(*P*) ∩ All(*Q*^∗^) ⊆ All(*P*) – so that, united with All(*P*), we obtain All(*P*). ◂

This leads to the following simplification – once again, the decrease in size of the query is immediate.

#### ▸ Corollary 10.

(𝒜, ℬ, 𝒞, 𝒟) *simplifies into* ({*A*^∪^}, ℬ^⋆^, 𝒞^⋆^, {*D*^∪^}), *where* ℬ^⋆^ = {*B* ∈ ℬ^∗^ :∄*B*^*′*^ ∈ ℬ^∗^, *B*^*′*^ ⊆ *B*} *and* 𝒞^⋆^ = {*C* ∈ 𝒞^∗^ : ∄*C*^*′*^ ∈ 𝒞^∗^, *C*^*′*^ ⊆ *C*}.

Note that this result also eliminates any duplicates in ℬ and 𝒞.

To identify these inclusions in practice, a straightforward algorithm would be to test for the inclusion of every possible pair of sets, with complexity *O*·(*η*(|ℬ^∗^|^2^ + |𝒞^∗^|^2^); where 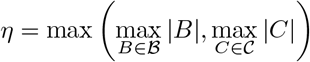 would be the size of the longest tag in either ℬ or 𝒞. It appears to be impossible to beat this worst-case complexity, unless the Strong Exponential Time Hypothesis [6] is false – see for instance this discussion on Theoretical Computer Science Stack Exchange [2].

However, in practice it can be useful to partition sets intelligently in order to limit the number of inclusion tests to be performed. With Algorithm 1 we propose such a method, running in *O*(*η* · |𝒫|^2^) – as per the previous discussion – provided that checking whether *f* (*n*) is defined for some *n* can be done in *O*(1).

#### Algorithm 1

RemoveInclusions

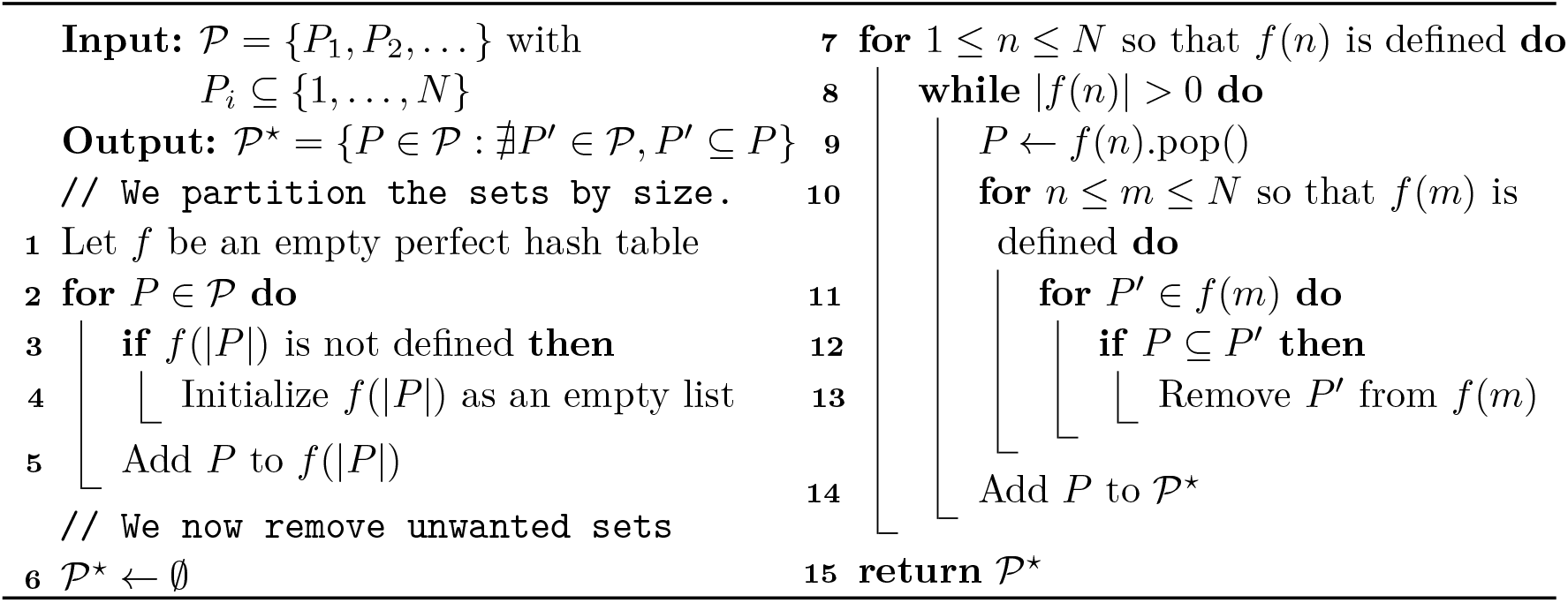

### 3.4 Simplification algorithm

Algorithm 2 summarises the results of this section.

#### Algorithm 2

QuerySimplifiation

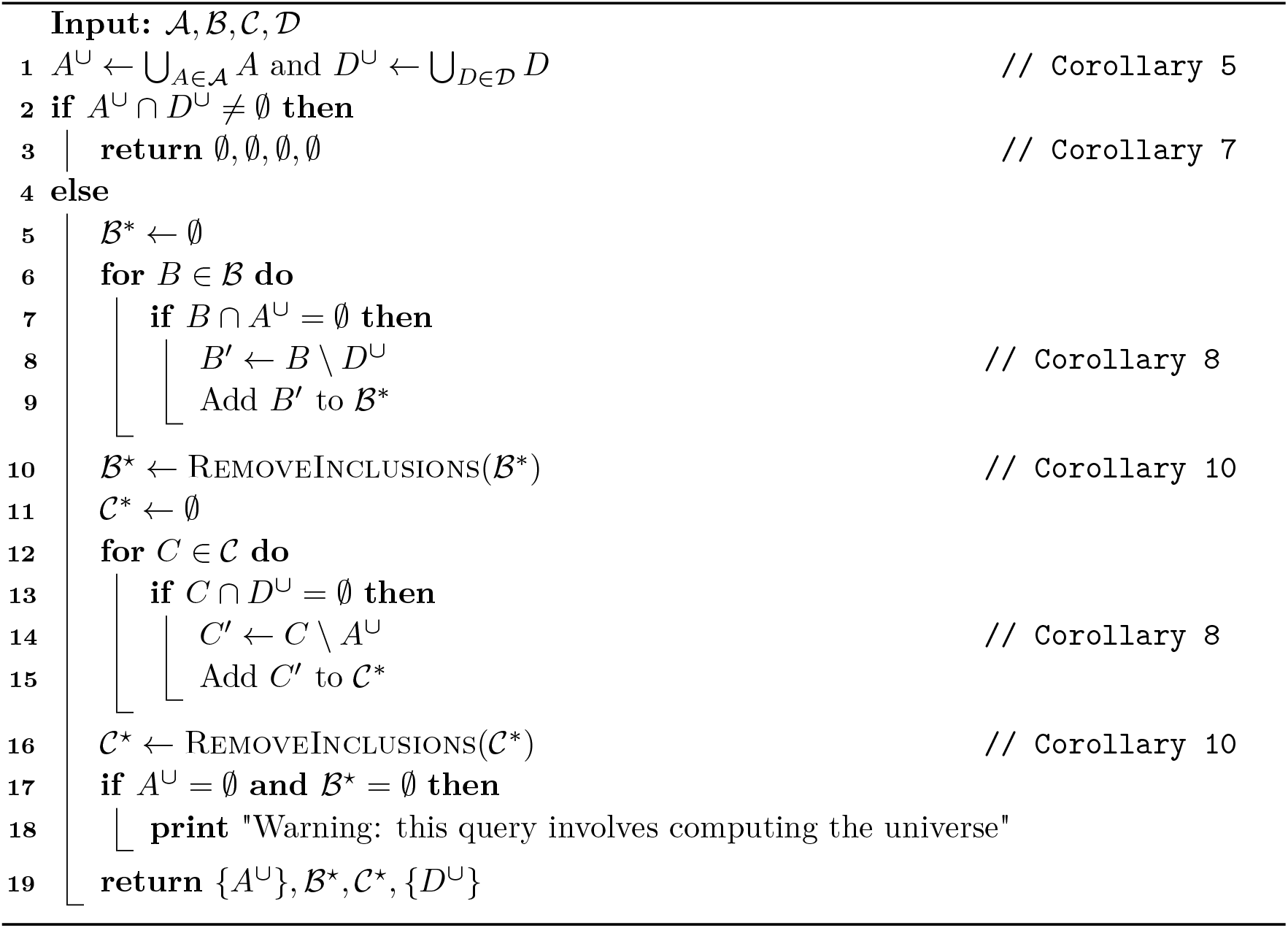

Note that it is possible to save some time by adding the sets *B*^*′*^ in line 9 (respectively *C*^*′*^ in line 15) not to ℬ^∗^ (respectively 𝒞^∗^) but rather directly to a hash table according to their size (as per lines 2–5 of Algorithm 1) before pursuing directly with the search for inclusions (lines 7–14 of Algorithm 1). Besides, and although we have not included it for the sake of simplicity, empty sets *B*^*′*^ or *C*^*′*^ are to be removed. In our implementation, Algorithm 2 is coded in Python, as its complexity relative to the task ahead of us – computing *σ*(𝒜, ℬ, 𝒞, 𝒟)– is not critical from a performance point of view.

#### ▸ Proposition 11.

*Algorithm 2 runs in*

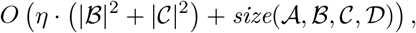

*where* 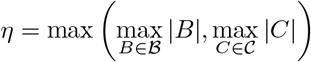.

**Proof**. Computing *A*^∪^ and *D*^∪^ is done in *O* (Σ_*A*∈𝒜_ |*A*| + Σ_*D*∈𝒟_ |*D*|) – using a hash table to represent sets, as done in Python with the type `set`.

Checking whether *B* ∩ *A*^∪^ = ∅ (respectively *C* ∩ *D*^∪^ = ∅) can be done in *O*(|*B*|) (respectively *O*(|*C*|)), as well as computing *B \ D*^∪^ (respectively *C \ A*^∪^). Summing over all *B* ∈ ℬ and all *C* ∈ 𝒞 lead the result, with the remainder that RemoveInclusions(𝒫) runs in *O*(*η* · |𝒫|^2^). ◂

## 4 Computation of *σ*(𝒜, ℬ, 𝒞, 𝒟)

After simplifying the query, we are left to compute the following quantity:

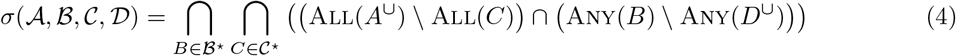

### 4.1 General strategy

For each 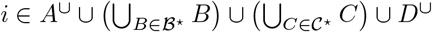 – that is, for each index required at some point in the query, we must access to *σ*(*R*_*i*_). Two options are available:

▄ Either the references *R*_*i*_ are stored in a compressed form that does not allow direct access to their set of *k*-mers, in which case *σ*(*R*_*i*_) must be constructed as a pre-processing step,
▄ or the references *R*_*i*_ are already stored in the form of a set of *k*-mers and all that is needed is to load *σ*(*R*_*i*_) into memory.

In the sequel, we use the Rust library CBL [11], which implements dynamic sets of *k*-mers efficiently, using a quotiented representation of *k*-mers. CBL supports fast insertion/deletion and set operations (union, intersection and difference), including in-place set operations and union/intersection of multiple sets in a single operation.

Our ambition is to restrict the use of RAM as much as possible, in a conservative fashion. To do this, no more than two CBL sets are manipulated in memory at the same time: the set being constructed, and the reference *σ*(*R*_*i*_) being processed at a given time. However, if one can afford to load several references into memory at once, certain steps described below (using union or intersection) can be performed in batches, thereby speeding up calculation time, thanks to the CBL library.

In addition, we prefer intersection operations (which can be carried out in place, and necessarily reduce the size of sets) over union operations (which involve copies of *k*-mers and increase the size of sets), whenever possible.

### 4.2 Initialization

This step has two possible versions, depending on whether *A*^∪^ = ∅ or not. In both cases, the aim is to construct an initial running set *S*, which will then be reduced in the subsequent construction.

**Case** *A*^∪^ *=* ∅ In this case, we start by constructing the set All(*A*^∪^) using the following procedure.

Let *a* ∈ *A*^∪^; load into memory *σ*(*R*_*a*_) – the initial running set. Then, for each *a*^*′*^ ∈ *A*^∪^, *a*^*′*^ ≠ *a*, load into memory *σ*(*R*_*a*_*′*) and intersect it with the current running set. This operation can be performed by batch, if applicable. At the end, we have built the set *S* = All(*A*^∪^).

**Case** *A*^∪^ = ∅ Again, this situation raises two alternatives.

▄ If ℬ^⋆^ ≠ ∅. Our initial running set will be one of the sets Any(*B*) for some *B* ∈ ℬ^⋆^. It seems only natural to construct the one that requires the fewest operations. Let *B* ∈ ℬ^⋆^ so that |*B*| is minimal. We have *S* = Any(*B*) = ∪_*b*∈*B*_ *σ*(*R*_*b*_). After constructing *S*, we remove *B* from ℬ^⋆^.
▄ otherwise, ℬ^⋆^ = ∅; we have no choice but to build the universe, i.e. 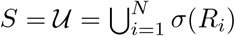. Notice that 𝒰 = Any({1, …, *N*}) – so we retrieve the previous case with an artificial *B* = {1, …, *N*}.

Eitherway, the running set is built by iteratively loading into memory each appropriated reference (with or without batches) and take the union with the previous ones – in a similar fashion as the computation of All(*A*^∪^) indicated above.

### 4.3 Treating 𝒞^⋆^

Let *S* be the running set at the end of the previous step. From (4), we now compute 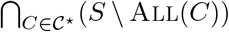. If 𝒞^⋆^ = ∅, skip this section. If not, the next procedure is to be followed. Note that for any sets *P* and *Q, P \ Q* = *P \* (*Q* ∩ *P*). Let us fix *C* ∈ 𝒞^⋆^. Instead of building All(*C*), we build instead *S* ∩ All(*C*). Let *S*^*′*^ be a copy of *S* – *S*^*′*^ is our *local* running set, whereas *S* is the *global* running set. Iteratively, we load into memory *σ*(*R*_*c*_) for each *c* ∈ *C*, and intersect it against the local running set. This part can be done with batches. At the end, the local running set is *S*^*′*^ = *S* ∩ All(*C*), and we replace *S* – the global running set – by *S \ S*^*′*^. We repeat this whole process for each remaining *C*^*′*^ ∈ *𝒞*^⋆^, *C*^*′*^*≠ C*.

### 4.4 Treating *D*^∪^

Let *S* be the running set at the end of the previous steps. We rewrite (4), as:

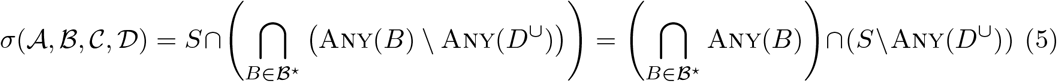

We are interested here in the term *S \* Any(*D*^∪^); we have

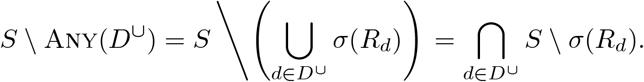

Noticing that(*P \ Q*) ∩ (*P\ R*) = (*P \ Q*) \ *R*, for any sets *P, Q* and *R*, we can compute *S \* Any(*D*^∪^) by successively replacing *S* by *S \ σ*(*R*_*d*_) for each *d* ∈ *D*^∪^.

Obviously, if *D*^∪^ = ∅, this step is skipped.

### 4.5 Treating ℬ^⋆^

Let *S* be the running set at the end of the previous steps. If ℬ^⋆^ = ∅, skip this step. Otherwise, from (5), we are left with:

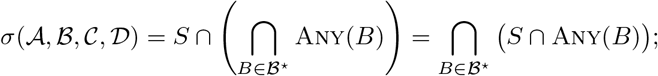

that can be sequentially computed as(((*S* ∩ Any(*B*_1_)) ∩ Any(*B*_2_)) ∩ …). Notice that

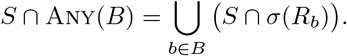

There is no escaping the calculation of an union here; we are again using a local running set *S*^*′*^ for this task. Let us fix some *B* ∈ ℬ^⋆^ and initialize the local set *S*^*′*^ as an empty set. For each reference *b* ∈ *B*, we compute *S* ∩ *σ*(*R*_*b*_) – notice that *S*, the global set, must not be modified here – and take the union of this set with *S*^*′*^. At the end of this process, replace *S* by *S*^*′*^ and repeat for each remaining *B*^*′*^ ∈ ℬ^⋆^, *B*^*′*^ ≠ *B*.

#### ▸ Remark 12.

Note that we adopt this procedure, rather than simply calculating *S* ∩ Any(*B*) directly, in order to limit the size of the sets manipulated. This is because the set Any(*B*) can be potentially much larger than *S*, while the set *S* ∩ *σ*(*R*_*b*_) is necessarily contained in *S*.

After dealing with ℬ^⋆^, the set *S* is exactly equal to *σ*(𝒜, ℬ, 𝒞, 𝒟).

### 4.6 Algorithm

Algorithm 3 sums up the procedure of this section. As a remainder, lower case letters (e.g. *b*) designate indices of *references*; capital letters (e.g. ℬ) designate sets of *indices*; calligraphic letters (e.g. *B*) designate sets of *sets of indices. S*_*l*_ designates the *local* running set – specific to the current step – whereas *S*_*g*_ designates the *global* running set, i.e. the solution set under construction.

#### Algorithm 3

QueryComputation

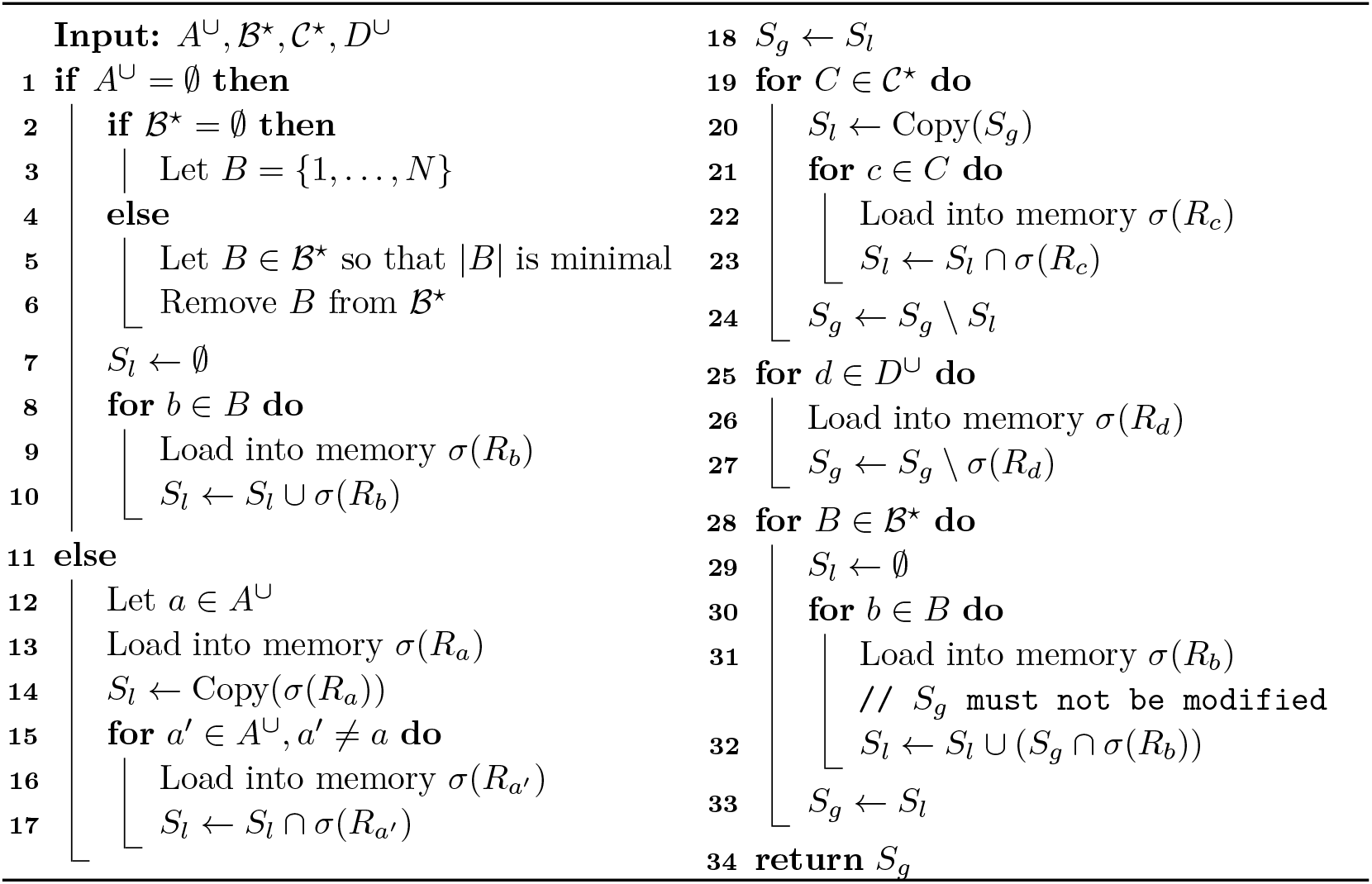

In order to analyse the complexity of Algorithm 3, we make the following assumptions.

i. We assume that the references are all of the same size, which we denote *K* – so that, ∀*i, K* = |*σ*(*R*_*i*_)|. We assume that a reference is loaded into memory in *O*(*K*) time;
ii. for any two *k*-mers sets *P* and *Q*, we assume that computing *P* ∩ *Q* can be done in *O*(min(|*P* |, |*Q*|)); that computing *P\Q* can be done in *O*(|*P* |); and that computing *P* ← *P* ∪ *Q* – i.e., extending *P* with the content of *Q* – can be done in *O*(|*Q*|).

#### ▸ Remark 13.

The complexities presented in (ii) are simplifications of the actual complexities of the operations as implemented in the CBL library. The interested reader can find their detailed complexity in Supplementary, Section A.2.

#### ▸ Proposition 14.

*Algorithm 3 admits the following complexities:*

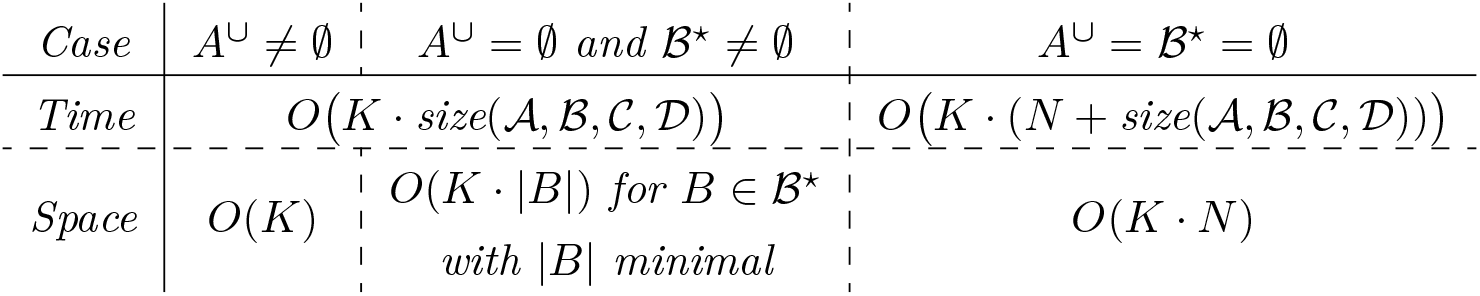

*where size*(𝒜, ℬ, 𝒞, 𝒟) *is defined in* (2).

Proof. The proof can be found in Supplementary, Section A.3. ◂

Remember that in terms of space, our approach is conservative; if one can afford to have several references loaded in memory, one can perform the unions or intersections in batches, which are faster in practice with the CBL library, as already stated earlier.

It is also clear that Section 3, designed to simplify the query, is of great interest in view of the time complexity of Algorithm 3.

## 5 Implementation

The implementation of our solution is composed of two main parts, a query builder in Python, and a wrapper for the CBL software written in Rust. CBL is a dynamic set structure for *k*-mers. The query builder reworks the initial query to minimize the number of operations and of files to load. The wrapper deals with fasta I/Os and executes Algorithm 3 according to the query refined by the query builder (implementing Algorithm 2). It benefits from recent advancements in CBL allowing set operations on multiple entries (batch). We provide a batch parameter *b* controlling the number of CBL sets loaded at the same time. This parameter is equal to 1 by default to limit memory usage. Choosing a larger *b* accelerates the computation of unions and intersections (since CBL can compute the union/intersection of *b* sets in a single operation) but increases peak memory usage.

Our implementation can be found at https://github.com/kamimrcht/Grimr.

## 6 Results

In the following, we test the most time consuming operations in the pipepline: set operations used in Algorithm 3. CBL allows to perform these operations in batches of size *X*, which means that it will apply a set operation simultaneously on *X* datasets. We evaluate the benefits of an increasing batch size in Subsection 6.1. We then show examples of our tool’s usage in Subsection 6.2. All experiments were performed on a single cluster node running with Intel(R) Xeon(R) Gold 6130 CPU @ 2.10GHz with 128GB of RAM and Ubuntu 22.04.

### 6.1 Batch size strategy for set operations

We analyse the execution time and *k*-mer processing rates at various batch sizes. This analysis aims to understand how batch processing impacts the performance of these operations on large genomic datasets. For each operation, we varied the batch size from 1 to 20 on 1000 bacterial Refseq genomes (1.7 billion of distinct *k*-mers) to assess the impact on performance (accessions in Supplementary, Section A.5).

We recorded the time taken for each batch size and calculated the *k*-mer processing rates. Figure 1 (top) shows that all operations shows a decreasing trend in execution time as the batch size increases. The input and output *k*-mers per second increase with batch size, indicating improved efficiency with larger batches. The peak RAM during those operations was 103 GB, when unioning all datasets. It is noteworthy that such an amount is not a reflect of most queries, and can in practice be an order of magnitude lower at least (an example is provided in Section 6.2 on a similar amount of *k*-mers).

**Figure 1.**
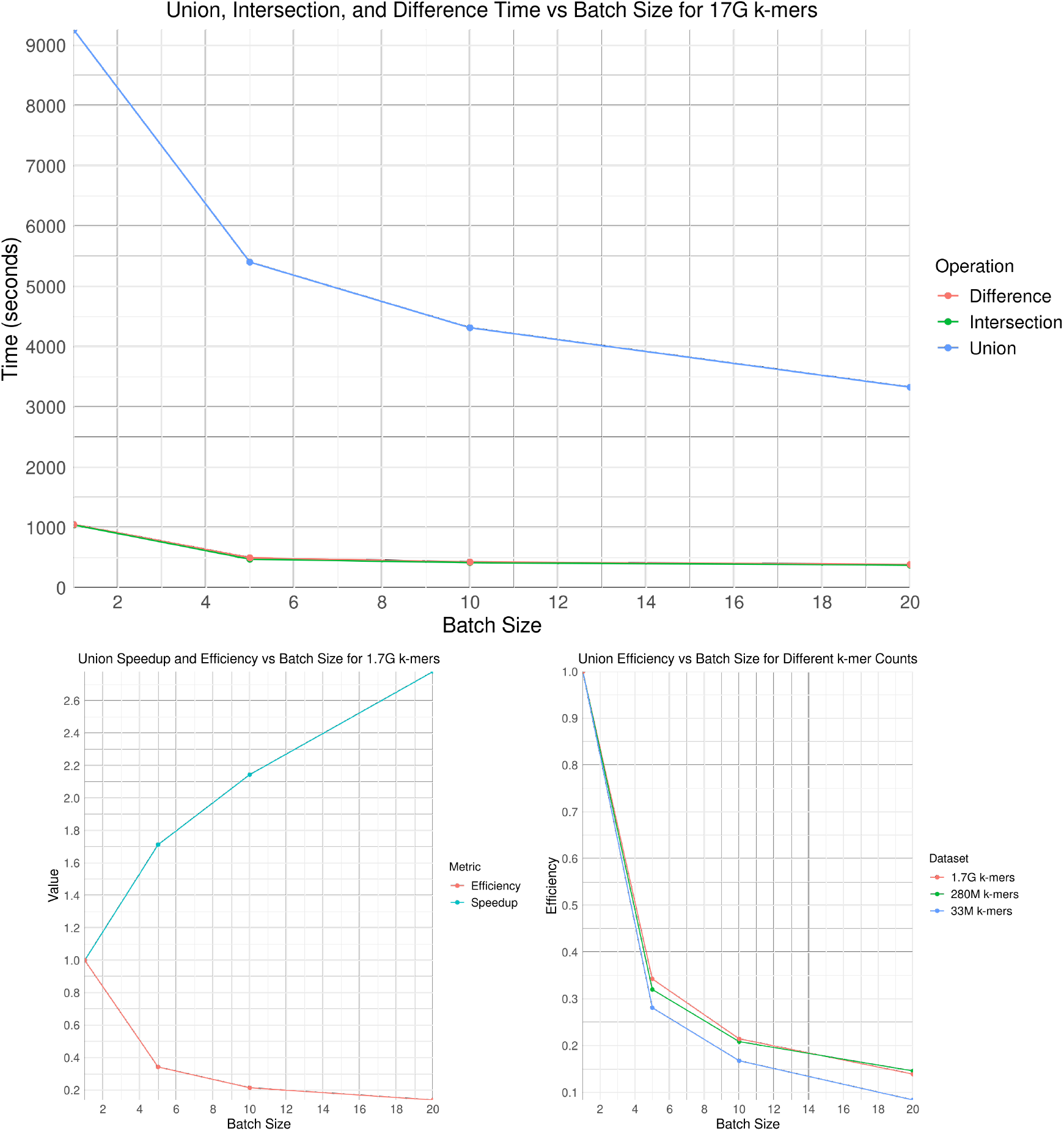
(Top) Time to realise union, intersection and difference for different batch sizes on 1.7G *k*-mers (1000 datasets). (Bottom left) Speedup and efficiency of the union operation on 1.7G of distinct *k*-mers from diverse bacterial genomes with increasing batch number. (Bottom right) Comparative efficiency for the union operation with increasing batch numbers, for different collections of samples (10 samples with 33M *k*-mers, 100 samples with 257M *k*-mers, 1000 samples with 1.7G *k*-mers).

We also inspected speedup and efficiency of the batch strategy. Speedup is defined as the ratio 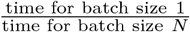 and efficiency as the ratio 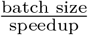. Here we report results for the union and observe similar behaviours for the other operations. Figure 1 (bottom left) shows that the speedup increases as the batch size increases, indicating that larger batch sizes reduce the time required for processing. However, the efficiency decreases as the batch size increases, suggesting that the gains have a diminishing return with larger batch sizes.

We also looked at the impact of batch effect’s scalability with an increasing number of *k*-mers using collections of 10, 100 and 1000 datasets (33 million, 257 million and 1.7 billion of distinct *k*-mers respectively). We focused on the union as it is our most costly operation. Figure 1 (bottom right) shows that the largest (1.7G and 257M) *k*-mers dataset generally shows better efficiency compared to the 33M *k*-mers dataset, particularly at higher batch sizes.

In order to assess similar results on different types of data, we produced similar experiments on RNA-seq human samples. For each operation, we varied the batch size from 1 to 20 on 200 human RNA-seq samples, up to 700 million of *k*-mers. The results present similar trends and are presented in Supplementary Figure S3.

### 6.2 Application cases

In the following, we tested the full piece of software in order to obtain subsets of *k*-mers relative to a query. We illustrate the query types in Figure 2 (a). We report the resources used for the runs, and illustrate excerpts of the results.

**Figure 2.**
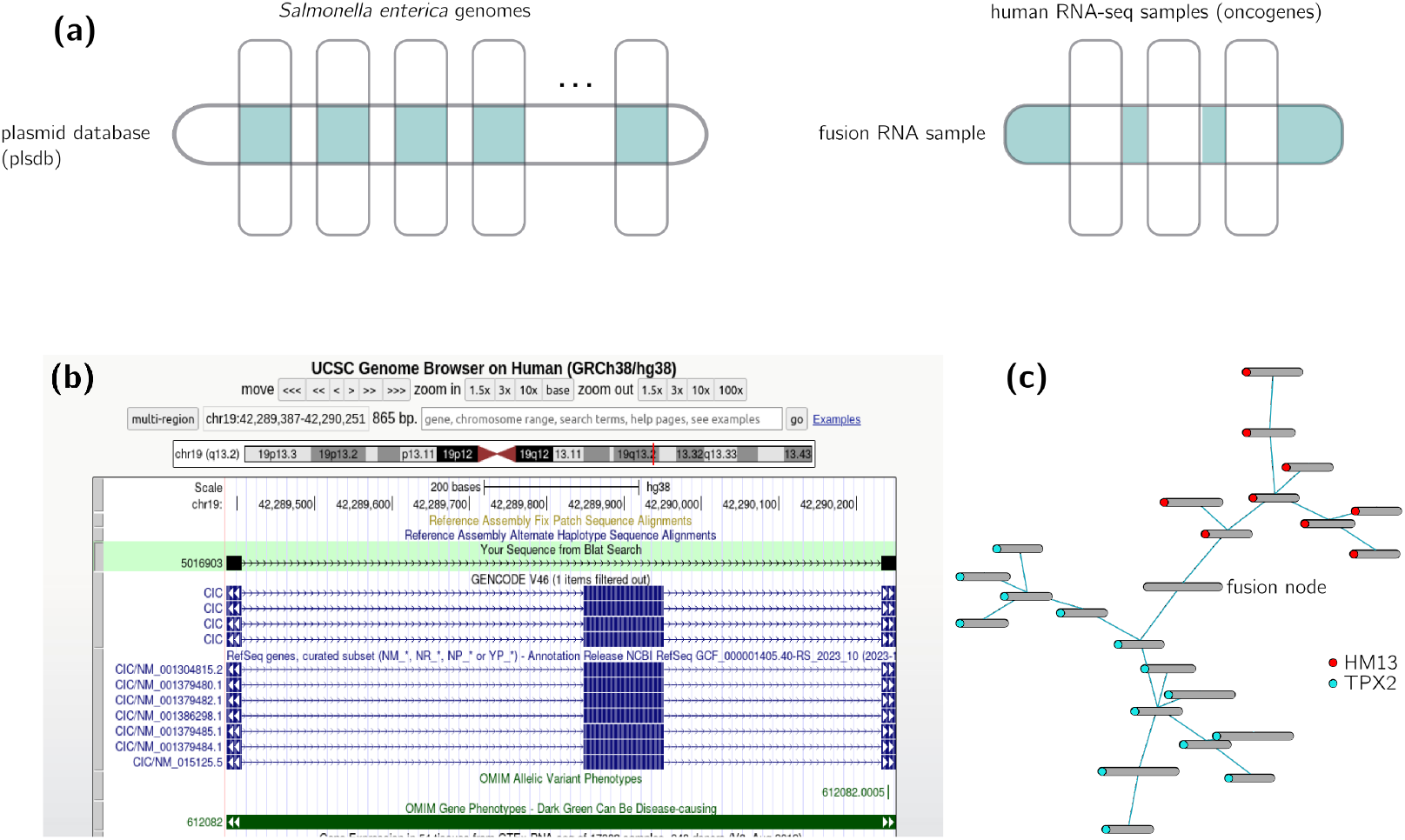
(a) Two types of queries demonstrated for the application cases. (b) *k*-mers exclusive to the fusion sample can lead to aberrant splicing. For instance, this sequence (track in green) from our findings mapped to the human genome (GRCh38) shows a neat un-annotated exon splicing in the CIC gene (annotated transcripts in the lower tracks). The figure was obtained using UCSC’s blat (https://genome.ucsc.edu/cgi-bin/hgBlat). (c) Another aberrant transcript. Another sequence in the exclusive subset from the fusion sample shows *k*-mers of a fusion transcript linking two genes (HM13 and TPX2). Since the mapped event spans a very long region of human chromosome 20, we chose to show the annotated de Bruijn graph locally. The fusion node contains *k*-mers yielded by our query, that clearly separates two subgraphs, each associated with nodes coming from a different gene (in red and blue). The figure was obtained using the de Bruijn graph visualisation tool Vizitig (https://gitlab.inria.fr/pydisk/examples/vizitig).

#### Find plasmid *k*-mers in bacterial genomes

We indexed the whole plasmid database (PLSDB [13]), as well as ten *Salmonella* genomes (accessions in Supplementary Table S1) in 1h15 min. We then queried 21-mers present at the intersection of the PLSDB with any of the genomes. The operation took 5 minutes and yielded 149,180 *k*-mers out of the 1.7 billion *k*-mers of PLSDB (75G of RAM).

#### Find *k*-mers unique to an experiment

We indexed several RNA-seq short reads datasets associated with putative oncogenes, including a *fusion_dataset* known for containing transcript fusion events (accessions in Supplementary Table S1), representing 74.5 million *k*-mers, in 5 minutes (3GB RAM). We queried the 21-mers exclusive to *fusion_dataset* in 51 seconds. We then assembled these *k*-mers using bcalm2 [3] (options: -kmer-size 21 -abundance-min 2), yielding ≈ 1.5 million of contigs, among which we obtained 134 contigs larger than 1,000 nucleotides. Manual exploration showed that *k*-mers from the results indeed displayed different aberrant splicing patterns, we show two examples in Figure 2 (b) and (c), illustrating the capacity of our method to identify *k*-mers of interest.

## 7 Conclusion

In this article, we presented a framework for the efficient enumeration of *k*-mers within a dataset that satisfy certain constraints on their metadata tags. Firstly, we proposed a user-friendly way of constructing queries, close to natural language, by freely associating tags with operations (All, Any, NotAll and NotAny). This query is then simplified in order to reduce computing delays. Secondly, the solution set of the query is constructed using CBL, a Rust library dedicated to optimised computation of set operations involving *k*-mers. Particular care has been taken to ensure that this construction is parsimonious both in terms of memory and computing time, in particular by operating in place whenever possible, in order to reduce unnecessary copies. An option is provided for batch processing of the data, should memory not be a limiting factor.

Our results demonstrate that larger batch sizes and datasets enhance performance and scalability, with unions up to 1.7 billion *k*-mers being feasible on a single cluster node. Practical applications, such as identifying plasmid *k*-mers in bacterial genomes and isolating unique *k*-mers in RNA-seq datasets, highlight the tool’s utility in genomic research.

This work allows a new type of query on genomic indexes, which we hope will lead to the discovery of new *k*-mers relevant to understanding biological phenomena, such as in the medical field where certain diagnoses rely on the detection of known patterns in the genome – and where the discovery of new patterns is therefore of utmost interest, as those that populate molecular signature databases [8].

Indexes capable of taking abundance into account are an ongoing major challenge, which has seen significant contributions in recent years, see for example [10, 7, 4]. The method we propose in this article is based solely on the presence/absence of *k*-mers; an interesting prospect for future work would be to take into account abundances – i.e. the number of times a *k*-mer appears in a reference – to allow an even wider range of queries.

## A Supplementary material

### A.1 Proof of Proposition 2

**Proof of (i), (ii) and (iii)** Let *x* ∈ All(*Q*). Then ∀*i* ∈ *Q, x* ∈ *σ*(*R*_*i*_). In particular, for any *j* ∈ *P* ∩ *Q, x* ∈ *R*_*j*_. Since *j* ∈ *P*, *x* ∈ Any(*P*). Then, All(*Q*) ⊆ Any(*P*). For any two sets *A* and *B*, we have *A* ⊆ *B* ⇐⇒ *A* ∪ *B* = *B* ⇐⇒ *A* ∩ *B* = *A* ⇐⇒ *A\ B* = ∅.

**Proof of (iv)** Let us denote *A* = All(*Q*^∗^), *B* = All(*P* ^∗^) and *C* = All(*I*). Then,

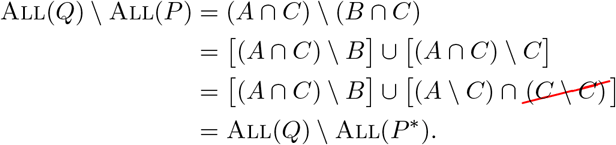

**Proof of (v)** Let us denote *A* = Any(*Q*^∗^), *B* = Any(*P* ^∗^) and *C* = Any(*I*). Then,

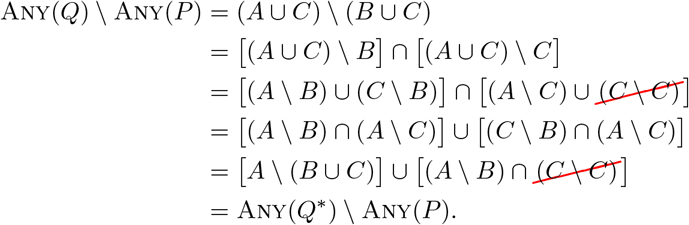

### A.2 Simplified complexity model for CBL

Internally, CBL represents *k*-mers as their smallest cyclic rotations, also known as necklaces, and stores them in a quotiented data structure: if two *k*-mers share the same necklace prefix of size *p*, this prefix is written only once and the suffixes are grouped together in bucket. The full description of this construction can be found in [11].

Provided a CBL set *A*, we denote its set of necklace prefixes as *P*_*A*_. Provided with a prefix *p* ∈ *P*_*A*_, we denote by *A*_*p*_ the set of *k*-mers of *A* whose necklace starts with *p*.

Every set operation between two sets *A* and *B* relies on first comparing *P*_*A*_ to *P*_*B*_ and then comparing the suffix buckets corresponding to prefixes in *P*_*A*_ ∩ *P*_*B*_. Thus, the complexity of any set operation is lower bounded by |*P*_*A*_| + |*P*_*B*_|.

Fortunately, since the necklace transformation preserves locality, |*P*_*A*_| is very small compared to |*A*|, usually 1-10% of |*A*|. What’s more, the prefixes are stored in a bitvector, so we can compare them very quickly, 64 comparisons at once.

To simplify our calculations, we assume that each bucket can be copied in constant time.

#### Union in place

When the union is done in place, i.e. *A* ← *A* ∪ *B*, we only need to iterate over the *k*-mers of *B*, which lowers the complexity:

▄ for each *p* ∈ *P*_*A*_ ∩ *P*_*B*_, we update *A*_*p*_ ← *A*_*p*_ ∪ *B*_*p*_ in *O*(|*B*_*p*_|).
▄ for each *p* ∈ *P*_*B*_ *\ P*_*A*_, *B*_*p*_ is copied in *O*(1).

This gives a total complexity of

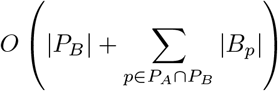

which is upper bounded by *O*(|*B*|).

#### Intersection

The intersection of *A* and *B* is computed as follows:

▄ for each *p* ∈ *P*_*A*_ ∩ *P*_*B*_, we compute *A*_*p*_ ∩ *B*_*p*_ in *O*(min(|*A*_*p*_|, |*B*_*p*_|)).
▄ for each *p* ∈ *P*_*A*_Δ*P*_*B*_, the corresponding bucket is discarded.

This gives a total complexity of

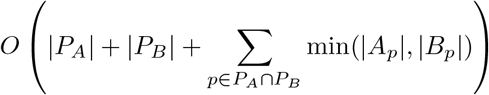

which is upper bounded by *O*(min(|*A*|, |*B*|) + max(|*P*_*A*_|, |*P*_*B*_|)).

#### Difference

The difference of *A* and *B* is computed as follows:

▄ for each *p* ∈ *P*_*A*_ ∩ *P*_*B*_, we compute *A*_*p*_ *\ B*_*p*_ in *O*(|*A*_*p*_|).
▄ for each *p* ∈ *P*_*A*_ *\ P*_*B*_, the corresponding bucket is copied in *O*(1).
▄ for each *p* ∈ *P*_*B*_ *\ P*_*A*_, the corresponding bucket is discarded.

This gives a total complexity of

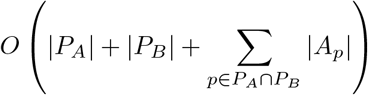

which is upper bounded by *O*(|*A*| + |*P*_*B*_|).

##### ▸ Remark 15.

For the proof of Proposition 2 (see next section), we made the assumptions that *A* ∩ *B* can be computed in *O*(min(|*A*|, |*B*|)); *A \ B* in *O*(|*A*|); and *A* ← *A* ∪ *B* in *O*(|*B*|).

As can be seen, these assumptions ignore the cost associated to prefixes – which are negligible in practice.

### A.3 Proof of Proposition 2

Complexities are analysed separately.

#### Space complexity

References require *O*(*K*) of space, and only one of them is loaded into memory at any given time. In addition, the solution set we construct is gradually reduced over the course of the algorithm, and is therefore maximal at initialisation – hence the result, depending on the case.

#### Time complexity

We start by analysing the conditional block from lines 1 to 17.

▄ If *A*^∪^ ≠∅, *S*_*l*_ is built in *O*(*K*). Then, for each *a*^*′*^ ∈ *A*^∪^, *a*^*′*^ ≠*a*, computing *S*_*l*_ ∩ *σ*(*R*_*a*_*′*) takes, as per assumption (ii), *O*(min(|*S*_*l*_|, |*σ*(*R*_*a*_*′*)|)), which is *O*(*K*). Summing over all *a*^*′*^ and taking into account *a* leads to a complexity of *O*(*K* · |*A*^∪^|);
▄ if *A*^∪^ = ∅ and ℬ^⋆^ ≠ ∅, assuming that acceding to |*B*| can be done in *O*(1), finding the smallest set *B* ∈ ℬ^⋆^ is done in *O*(|ℬ^⋆^|). By assumption (ii), building *S*_*l*_ in lines 8–10 is done in *O*(*K* · |*B*|);
▄ if *A*^∪^ = ℬ^⋆^ = ∅, building *S*_*l*_ in lines 8–10 is done in *O*(*K* · *N*) by assumption (iii).

For lines 19–24, assuming 𝒞^⋆^ ≠ ∅; let us fix *C* ∈ *𝒞*^⋆^. Building *S*_*l*_ in line 20 is *O*(*K*); the loop of lines 21–23 is *O*(*K* · |*C*|) – as per assumption (ii). Line 24 is *O*(|*S*_*g*_|), which is bounded by *O* (*K*). Therefore, summing over all *C* ∈ 𝒞 ^⋆^, we get 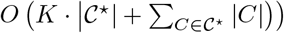, which can be simplified to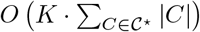.

Lines 25–27 are in *O*(*K* · |*D*^∪^|), again since |*S*_*g*_| ≤ *K* and with assumption (ii).

Finally, for lines 28–33, let us fix *B* ∈ *ℬ* ^⋆^. At line 32, computing *S*_*g*_ ∩ *σ*(*R*_*b*_) is *O*(min(|*S*_*g*_|, |*σ*(*R*_*b*_)|)) which is *O*(*K*); since |*S*_*g*_ ∩ *σ*(*R*_*b*_)| ≤ *K*, computing *S*_*l*_ ∪ (*S*_*g*_ ∩ *σ*(*R*_*b*_)) is also *O*(*K*). Summing over all *B* ∈ *ℬ* ^⋆^ leads to 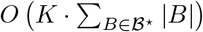. Note that if one set *B* was removed from ℬ^⋆^ at the initialisation (in the case *A*^∪^ = ∅ and ℬ^⋆^ ≠ ∅), summing the complexity of initialisation with the complexity of lines 28–33 does indeed lead to the same complexity, i.e. 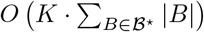.

In the end, combining all the intermediate complexities leads to the result.

### A.4 Supplementary results

**Figure S3.**
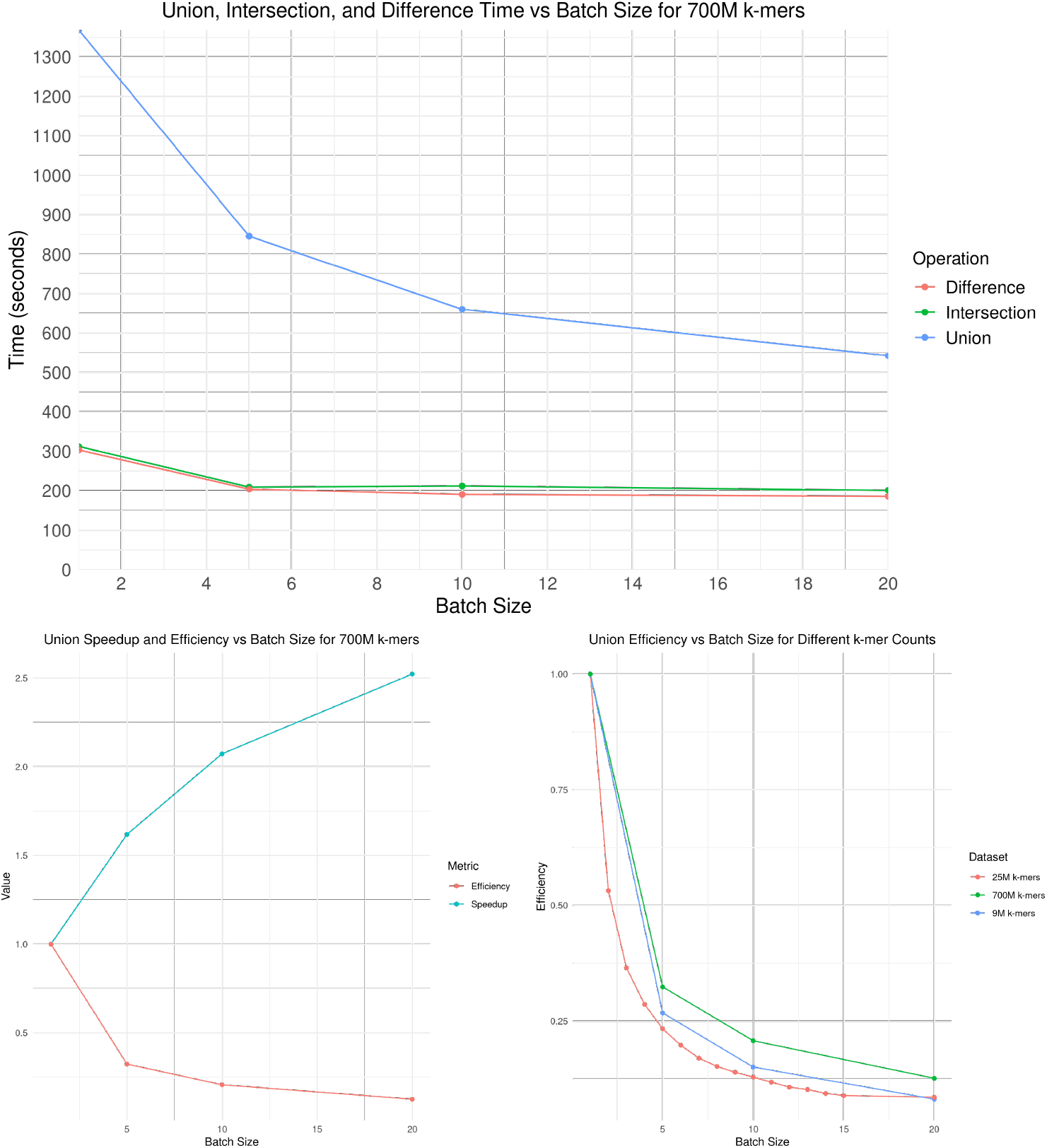
(Top) Time to realize union, intersection and difference for different batch sizes on 700M *k*-mers (100 datasets). (Bottom left) Speedup and efficiency of the union operation on 700M of *k*-mers from human RNA-seq samples with increasing batch number. (Bottom right) Comparative efficiency for the union operation with increasing batch numbers, for different collections of samples (50 samples with 9M *k*-mers, 100 samples with 25M *k*-mers, 200 samples with 700M *k*-mers).

### A.5 Accessions

In addition to Table S1, the full lists of accessions for the benchmark on 1000 bacterial genomes, as well as for the benchmark on 200 human RNA-seq, can be found at https://github.com/kamimrcht/Grimr/blob/main/accessions.txt.

**Table S1.** Summary of Accessions from GenBank and SRA for application cases.

| Species | Type | Bank | Accessions |
| --- | --- | --- | --- |
| Salmonella enterica | Genome | GenBank | GCA_011232695.1_PDT000673809.1_genomic.fna.gz |
|  |  |  | GCA_015098575.1_PDT000855768.1_genomic.fna.gz |
|  |  |  | GCA_007565205.1_PDT000297230.2_genomic.fna.gz |
|  |  |  | GCA_015034225.1_PDT000869352.1_genomic.fna.gz |
|  |  |  | GCA_005401825.1_PDT000491193.1_genomic.fna.gz |
|  |  |  | GCA_021279925.1_PDT001211007.1_genomic.fna.gz |
|  |  |  | GCA_011288685.1_PDT000626723.1_genomic.fna.gz |
|  |  |  | GCA_022434075.1_PDT001256364.1_genomic.fna.gz |
|  |  |  | GCA_008929265.1_PDT000602336.1_genomic.fna.gz |
|  |  |  | GCA_017486225.1_PDT000985998.1_genomic.fna.gz |
| Human | Reads RNA-seq | SRA | SRR8615240 |
|  |  |  | SRR8615242 |
|  |  |  | SRR8616107 |

## Notes

### Competing Interest Statement

The authors have declared no competing interest.

## References

1 Chloé Bessière, Haoliang Xue, Benoit Guibert, Anthony Boureux, Florence Rufflé, Julien Viot, Rayan Chikhi, Mikaël Salson, Camille Marchet, Thérèse Commes, and Daniel Gautheret. Exploring a large cancer cell line rna-sequencing dataset with k-mers. bioRxiv, 2024. doi:10.1101/2024.02.27.581927.

2 Andreas Björklund. What is the fastest way to check for set inclusion? Theoretical Computer Science Stack Exchange. URL: https://cstheory.stackexchange.com/q/9897.

3 Rayan Chikhi, Antoine Limasset, and Paul Medvedev. Compacting de bruijn graphs from sequencing data quickly and in low memory. Bioinformatics, 32(12):i201–i208, 2016.

4 Mitra Darvish, Enrico Seiler, Svenja Mehringer, René Rahn and Knut Reinert. Needle: a fast and space-efficient prefilter for estimating the quantification of very large collections of expression experiments. Bioinformatics, 38(17):4100–4108, 2022.

5 Martin Hunt, Leandro Lima, Wei Shen, John Lees, and Zamin Iqbal. Allthebacteria - all bacterial genomes assembled, available and searchable. bioRxiv, 2024. doi:10.1101/2024.03.08.584059.

6 Russell Impagliazzo and Ramamohan Paturi. On the complexity of k-sat. Journal of Computer and System Sciences, 62(2):367–375, 2001.

7 Mikhail Karasikov, Harun Mustafa, Daniel Danciu, Marc Zimmermann, Christopher Barber, Gunnar Rätsch, and André Kahles. Indexing all life’s known biological sequences. bioRxiv, 2024. doi:10.1101/2020.10.01.322164.

8 Arthur Liberzon, Aravind Subramanian, Reid Pinchback, Helga Thorvaldsdóttir, Pablo Tamayo, and Jill P Mesirov. Molecular signatures database (msigdb) 3.0. Bioinformatics, 27(12):1739–1740, 2011.

9 Camille Marchet, Christina Boucher, Simon J Puglisi, Paul Medvedev, Mikaël Salson, and Rayan Chikhi. Data structures based on k-mers for querying large collections of sequencing data sets. Genome research, 31(1):1–12, 2021.

10 Camille Marchet, Zamin Iqbal, Daniel Gautheret, Mikaël Salson, and Rayan Chikhi. Reindeer: efficient indexing of k-mer presence and abundance in sequencing datasets. Bioinformatics, 36(Supplement_1):i177–i185, 2020.

11 Igor Martayan, Bastien Cazaux, Antoine Limasset, and Camille Marchet. Conway-Bromage-Lyndon (CBL): an exact, dynamic representation of k-mer sets. In 32nd International Conference on Intelligent Systems for Molecular Biology (ISMB 2024), 2024. doi:10.1101/2024.01.29.577700.

12 Amatur Rahman and Paul Medvedev. Representation of k-mer sets using spectrum-preserving string sets. In International Conference on Research in Computational Molecular Biology, pages 152–168. Springer, 2020.

13 Georges P Schmartz, Anna Hartung, Pascal Hirsch, Fabian Kern, Tobias Fehlmann, Rolf Müller, and Andreas Keller. Plsdb: advancing a comprehensive database of bacterial plasmids. Nucleic Acids Research, 50(D1):D273–D278, 2022.

